# Single neurons in the human medial temporal lobe flexibly shift representations across spatial and memory tasks

**DOI:** 10.1101/2023.02.22.529437

**Authors:** Thomas Donoghue, Runnan Cao, Claire Z Han, Cameron M Holman, Nicholas J Brandmeir, Shuo Wang, Joshua Jacobs

## Abstract

Investigations into how individual neurons encode behavioral variables of interest have revealed specific representations in single neurons, such as place and object cells, as well as a wide range of cells with conjunctive encodings or mixed selectivity. However, as most experiments examine neural activity within individual tasks, it is currently unclear if and how neural representations change across different task contexts. Within this discussion, the medial temporal lobe is particularly salient, as it is known to be important for multiple behaviors including spatial navigation and memory, however the relationship between these functions is currently unclear. Here, to investigate how representations in single neurons vary across different task contexts in the MTL, we collected and analyzed single-neuron activity from human participants as they completed a paired-task session consisting of a passive-viewing visual working memory and a spatial navigation and memory task. Five patients contributed 22 paired-task sessions, which were spike sorted together to allow for the same putative single neurons to be compared between the different tasks. Within each task, we replicated concept-related activations in the working memory task, as well as target-location and serial-position responsive cells in the navigation task. When comparing neuronal activity between tasks, we first established that a significant number of neurons maintained the same kind of representation, responding to stimuli presentations across tasks. Further, we found cells that changed the nature of their representation across tasks, including a significant number of cells that were stimulus responsive in the working memory task that responded to serial position in the spatial task. Overall, our results support a flexible encoding of multiple, distinct aspects of different tasks by single neurons in the human MTL, whereby some individual neurons change the nature of their feature coding between task contexts.

## Introduction

A key question in neuroscience concerns the nature of the information represented by the activity of individual neurons. A remarkable collection of studies have demonstrated how individual neurons can encode specific variables of interest at various levels of complexity, across perception, motor control, and cognitive tasks, ranging from orientation-tuned cells in primary visual cortex (Hubel & Wiesel, 1962), to abstract invariant representations of specific identities, such as the ‘Jennifer Anniston’ neuron (Quiroga et al., 2005). However, not all individual neurons encode with such specificity, with many experiments finding cells with conjunctive encoding, whereby an individual neuron responds to the combination of two or more features (Duvelle et al., 2023) and/or be even more broadly tuned to a mixture of multiple variables (Fusi et al., 2016; Rigotti et al., 2013). Collectively, this literature establishes that the brain contains a mixture of ‘specialist’ cells, which are narrowly tuned to a specific feature, as well as ‘generalist’ cells, which have broader mixed tuning across multiple variables, with many remaining open questions regarding if and when different encodings shift or remap entirely.

Apparent differences in the specificity of neural representations are at least partly related to regional differences. Where primary sensory areas are generally thought to be specifically tuned to incoming sensory information of specific modalities (though see a notable example of multi-sensory responses in V1 (Knöpfel et al., 2019)), increasingly ‘higher-order’ areas are thought to encode increasingly abstract constructs that could be more broadly tuned across tasks and contexts. The prefrontal cortex, for example, is thought to engage in high-level, abstract representations that can be flexibly applied across task contexts (Behrens et al., 2018; Duncan, 2001). Overall, there is thought to be a hierarchy of encoding from specialized neurons in primary sensory areas that are not expected to change their representation, to higher-order areas that encode more abstract features, with more flexibility to encode multiple features, both simultaneously and/or across time. Further understanding the consistency of representations, however, requires dedicated work that evaluates neural activity across task contexts.

In the cortex, some experiments have examined different task variants within a cognitive domain, to investigate how individual neurons change their activity in relation to different task demands. For example, in recordings in the parietal cortex from mice, neural responses across two different visual decision tasks engaged largely distinct populations of neurons (Lee et al., 2022). However, when using two different categorization tasks in monkeys, a broadly similar neural representation was found across the two tasks (Mohan et al., 2021). An early experiment in humans did report a small number of MTL neurons that showed responses to words and also to unrelated faces (Heit et al., 1988). Some studies in humans have examined responses to the same stimuli based on task demands, for example, finding differences in responses across regions to different task demands when viewing the same faces (Cao, Todorov, et al., 2022). In another experiment, asking participants to flexibly switch between a recognition memory and categorization task led to different population-level representations of the task demands in the medial frontal cortex (Minxha et al., 2020). Collectively, these studies are beginning to establish how neurons change their representations across variations in task context, however the limited results thus far suggests there are differences across species, cognitive domains, and anatomical locations.

Within the discussion of neural representations, the medial temporal lobe (MTL) is a structure of particular interest due to its involvement in multiple cognitive processes. In spatial navigation, the MTL has seemingly ‘specialist’ cells that encode specific locations in space (place cells), as well as location- and navigation-related features such as head direction, speed, and environment borders (Moser et al., 2017). While much of the spatial navigation literature is in rodents, space-related representations have also been found in non-human primates (Rolls & Wirth, 2018), as well as recent demonstrations of place, target-location, and sequence encodings in single-neurons in humans (Miller et al., 2013; Tsitsiklis et al., 2020). In investigations with visually presented stimuli, the MTL has also been found to have neurons that respond to broadly tuned object categories as well as to highly specific concepts, which is also thought to relate to memory processes (Quiroga, 2012; Rutishauser et al., 2021). Despite a large amount of research on both the spatial navigation and memory functions of the MTL, this research is typically done in distinct labs and experiments, with the relationship between memory and spatial-navigation functions of the MTL remains a debated topic. Specifically, it is unclear how the firing patterns of individual neurons shift between these different processes.

The previous findings, whereby the MTL has been found to engage in seemingly distinct functions across spatial navigation and memory has led to the suggestion that this structure may engage in representing ‘cognitive maps’ whereby individual neurons represent features or relations within a high-dimensional state space (Behrens et al., 2018; Schiller et al., 2015; Tolman, 1948). Under this hypothesis, the MTL constructs ‘maps’ of features of interest within which physical space may simply be a special case of a more general mechanism that can be applied to other ‘spaces’. In a physical context, the activity of place cells represent locations in the map, representing physical space. In a different context, individual MTL neurons are predicted to be able to represent elements within other feature spaces, such that they can flexibly engage in different kinds of representations based on task demands. This perspective is supported by studies such as one that finds ‘frequency-place cells’ wherein individual neurons represent locations in frequency space during a sound modulation task (Aronov et al., 2017). This framework is also consistent with perspectives whereby the hippocampus can be thought of as a general relational processing system which can be applied to organizing relations across space, time, and conceptual dimensions (Eichenbaum & Cohen, 2014).

Empirically, the specificity, flexibility, and consistency of neural responses can be examined by using different tasks while comparing the responses of a specific neural population across stimuli and task contexts. However, since the vast majority of experiments only investigate neural representations within specific behavioral contexts, it is generally unclear if and when individual neurons maintain and/or change representation across different contexts. Experiments employing multiple tasks are limited by practical challenges related to training and testing across multiple distinct tasks while recording from the same neurons, especially for tasks from across different cognitive domains. These limitations may be particularly difficult if attempting to train animal models to flexibly switch between completely distinct tasks. Such experimental designs are therefore a situation in which human participants may be ideal due to their capacity to rapidly learn and switch between distinct behavioral contexts.

Collectively then, the existing evidence suggests that the brain appears to use a hierarchically organized combination of ‘specialist’ and ‘generalist’ cells. However, beyond this general conceptualization, there are numerous outstanding questions relating to the nature of how individual neurons encode features across different task contexts. This question is particularly salient in the MTL, in which there is a strong theoretical motivation for predicting that individual cells engage in distinct representations across different contexts as part of domain-general ‘cognitive maps’. This is empirically well demonstrated when considering different contexts within the same domain (e.g. remapping across contexts in spatial experiments (Kubie et al., 2020; Kubie & Muller, 1991)), however there is currently little empirical investigation comparing across domains such as comparing across a spatial navigation task and a non-spatial visual memory task. To what extent do individual MTL neurons maintain a fixed encoding, responding only to a narrowly tuned feature structure across different behavioral contexts and disengaging if that feature is not present? Alternatively, can individual neurons be flexibly recruited to encode task-relevant variables, with potentially different representations based on behavioral context?

To address these questions, in this study, we investigate how the representations of individual neurons in the MTL change across task context. We do so using a paired-task design in human participants, while recording single-neuron activity from implanted electrodes. By comparing neural activity within and between distinct tasks, we hypothesized that we would observe individual neurons switch between distinct representations across different tasks. This is indeed what we observed, as we found both neurons that maintained a consistent representation to stimuli that were presented across tasks, as well as some neurons that engaged in seemingly distinct representations between the two tasks. This implies that some ‘specialist’ encodings may be specific to task context, and that (at least some) MTL neurons can switch the kind of feature that they encode, which may relate to the MTL as enacting a feature-general ‘cognitive map’ rather than being purely specialized within particular domains.

## Methods

### 2.1 Single-Neuron Recordings

The participants in our study were patients with chronic, medication-resistant epilepsy who volunteered to participate in our study while undergoing pre-surgical monitoring with implanted electrodes to localize epileptogenic regions. Patients were eligible for participation in this study if the clinical monitoring plan included electrode coverage in the medial temporal lobe (MTL), including unilateral or bilateral amygdala and/or hippocampus. Eligible participants were implanted with Behnke-Fried electrodes, which include clinical macro-electrodes as well as 40-μm microwires which extend from the macro electrode tip and can record single-neurons. Five patients (4 female, ages 29-53 years old) participated, for a total of 22 recording sessions (Table 1). Recordings were collected at J. W. Ruby Memorial Hospital, affiliated with West Virginia University, and all patients provided informed consent. Patients had between 3-6 Behnke-Fried electrodes implanted, to a total of 22 electrodes across the group, each of which had 8 micro-wires. Recordings from the microwires were collected at 32 kHz using a NeuraLynx Atlas recording system (Neuralynx, Bozeman, USA) with full bandwidth recordings (0.1-9 kHZ). The data analyzed in this study is openly available for the one-back task (Cao, Lin, et al., 2022) and will be released for the Treasure Hunt task upon publication.

**Table 1.** Subject overview. Details of the data that was included in this study.

| Subject ID | Age | Sex | Race | Electrode location | Number of sessions | Number of total units | Number of kept units |
| --- | --- | --- | --- | --- | --- | --- | --- |
| 001 | 29 | F | Caucasian | Bilateral amygdala & hippocampus | 4, 2 | 286, 229 | 245, 134 |
| 002 | 53 | F | Caucasian | Bilateral amygdala & hippocampus | 2, 3 | 156, 116 | 120, 95 |
| 003 | 49 | M | Caucasian | Bilateral amygdala & hippocampus | 1, 2 | 34, 86 | 18, 44 |
| 004 | 31 | F | Caucasian | Bilateral amygdala & hippocampus | 2, 4 | 262, 380 | 183, 264 |
| 005 | 26 | F | Caucasian | Left amygdala, bilateral anterior hippocampus & left posterior hippocampus | 1, 1 | 77, 45 | 41, 30 |
| Total |  |  |  |  | 10, 12 | 815, 856 | 608, 565 |

### 2.2 Experimental Tasks

Participants in our study performed a paired task session in which they completed a visual working memory task, which was immediately followed by a spatial navigation and episodic memory task. The visual working memory task was a one-back paradigm, which is commonly used to test how participants maintain and manipulate information in working memory (Cao et al., 2021; Cao, Wang, et al., 2022). The spatial navigation task is a 3D virtual stimulus-location associative memory task called Treasure Hunt (TH), developed in Unity, which was previously used to study various aspects of human spatial memory and electrophysiology (Miller et al., 2018; Tsitsiklis et al., 2020). Both tasks were played on bedside laptops, with participants pressing the spacebar on a keyboard to respond in the one-back and using a separate joystick to control movements in the Treasure Hunt task. In each paired-task session, participants typically started with the one-back task first and played Treasure Hunt immediately after, with short breaks between tasks if necessary.

Each of these tasks could occur in one of two versions, a “face” version where presented items were famous faces, or an “object” version where items were images of general objects. Within a single paired task session, the stimuli type was always consistent across tasks (eg. the object version of the one-back task was always followed by the object version of TH). In the “object” version of the one-back task, 10 images of objects from 50 categories were taken from the ImageNet database (Deng et al., 2009). This was paired with the standard version of Treasure Hunt in which stimulus items are 3d rendered images of everyday objects. In the faces version of the one-back, images of celebrities were taken from the CelebA dataset52 (Liu et al., 2015), selecting identities across genders and races. To create a “faces” version of treasure hunt, stimuli were replaced with a selection of face images that were also used in the one-back task. In the one-back task, for both versions of the tasks, we selected 50 identities and object categories with 10 images for each, resulting in 1000 images in total. The same set of stimuli were used across all patients.

We used a one-back task with human faces or objects as described in previous studies (Cao et al., 2021; Cao, Wang, et al., 2022). Stimuli were presented using MATLAB with the Psychtoolbox 3 (Kleiner et al., 2007) running a laptop computer with a screen resolution of 1600 × 1280. In each trial, a single image was presented at the center of the screen for 1 second. The interstimulus-interval (ISI) was uniformly sampled between 0.5 to 0.75 seconds. Patients were instructed to respond by pressing the spacebar if the current image was identical to the immediately preceding image. Patients were instructed to respond after the image disappeared, to avoid motor activity during image presentations. One-back repetition presentations happened on 9% of trials. Other than repetition trials, each individual image was presented once, with the order of presented images being randomized for each patient. All analyses were done excluding the one-back repetition trials, in order to have an equal number of responses for each image.

The Treasure Hunt spatial navigation and memory task was used, as described in previous studies (Miller et al., 2018; Tsitsiklis et al., 2020). In each trial of Treasure Hunt, participants use a joystick to navigate a rectangular arena on a virtual beach and encounter treasure chests that can contain items. Participants are instructed to remember the location of the presented items, so that they can later report the location of each encountered item. Chests appear in the arena one at a time with randomized locations across trials. There are typically 5 blocks of trials in a complete run of Treasure Hunt, where each block has 8 individual trials, to a total of 40 trials. Due to the time constraints of our paired-task session, most Treasure Hunt runs were not a complete session, with most runs instead including 3 complete blocks (24 trials). A total of 22 paired-task sessions were included and analyzed (TH-face: 10 sessions, average of 23.40 trials (range: 8-40); TH-object: 12 sessions, average of 25.67 trials (range: 24-32)).

Each trial of Treasure Hunt consists of two phases – a navigation / encoding phase, and a retrieval phase. During the navigation phase, participants are first placed at one end of the arena, from which they can navigate freely using the joystick. They are instructed to navigate to a series of chests that are presented serially in the arena. Upon reaching a chest, players are rotated to the front of the chest, at which point it opens and reveals either an item contained in the chest, which is presented for 1.5 seconds, or the chest is shown to be empty. There are 4 chest presentations per trial, 2 or 3 of which are full chests. After reaching all four chests of a trial, participants are transported to one end of the arena, either the same side as the navigation start or the opposite side, indicating the end of the navigation phase. Participants then play a distractor task which is a computerized version of the “shell game”. After the distractor game, the recall phase starts in which participants are prompted with each of the items from the trial in a random order. They are first asked to rate their confidence level of whether they remember the location of the encountered item (response options: “Yes”, “Maybe” or “No”). They are then asked to respond with the exact location in the arena they encountered the item by maneuvering a crosshair with the joystick. At the end of the recall period, participants receive feedback regarding whether each response is close enough to be considered correct, and receive points accordingly. A response is considered correct if it is within 13 virtual units of the true object location.

### 2.3 Data Pre-Processing

After data collection, each paired-task session was pre-processed together such that the same putative single-neurons could be isolated and analyzed across both tasks. Single-neuron activity was identified and spike sorted using the OSort algorithm (Rutishauser et al., 2006). Spike times were then extracted from the full session for each task, such that each task could be analyzed independently. Only neurons with an average firing rate greater than 0.15 Hz across both tasks were kept for analysis, keeping a total of 1173 neurons across the project (face-version: 608 neurons; object-version: 565 neurons). In the analysis, neurons from each recording session were considered unique, even if they came from the same subject. We note that the same participants did both stimulus versions of the paired-task sessions, such that there could be overlapping neurons in the two datasets. Notably, the two stimulus versions were analyzed independently. For the one-back task, single-neuron activity was associated with the behavioral timestamps and analyzed using custom scripts in the Matlab programming language (Mathworks, Inc, USA). For the Treasure Hunt task, single-neuron activity was organized together with behavioral information into Neurodata Without Borders (NWB) files (Rübel et al., 2022) which were then analyzed in the Python programming language, using the *spiketools* module for analyzing single-neuron activity (Donoghue et al., 2023). After each task was analyzed, the activity patterns and task activations within each task were then compared across tasks in order to examine single-neuron activity across task contexts.

### 2.4 Neural Data Analyses

Across both tasks, we analyzed neural responses that were potentially consistent across both tasks, such as responses to stimuli, and also responses that were specific to each task, for example, identity responses in the one-back task (in which multiple images of the same persons / objects are shown) and place and sequence-related responses in the Treasure Hunt task. To test for stimulus-related responses, we used t-tests comparing firing pre & post stimulus onset. To analyze cell responses to other task features, we used ANOVAs. For all ANOVA tests, we used surrogates to evaluate statistical significance, creating 1000 surrogates by circular shifting spike times by a random offset, recomputing the ANOVA of interest, and computing the f-statistics output from the real data as compared to the distribution of surrogates to compute an empirical p-value. For each analysis, individual neurons were considered significant at an alpha value of 0.05 (if the f-value calculated on the real data was at or above the 95th percentile of the f-value from the shuffled surrogate data). For all analyses, at the group level we applied one-sided binomial tests to evaluate whether the number of neurons detected to have a significant response exceeded the number expected by chance.

In the one-back task, we analyzed neurons for non-selective responses to stimuli, as well as for selective responses to specific stimuli, following previously described procedures (Cao et al., 2021; Cao, Wang, et al., 2022). To detect stimulus-responsive neurons, we used a paired *t*-test (p<0.05) comparing the firing rate during baseline (−250 ms to 0 relative to stimulus onset) versus stimulus period (250 ms to 1250 ms after stimulus onset). For the selection of identity neurons, we first used a one-way ANOVA (p < 0.05) to identify neurons that responded differently to different identities. In addition, we applied an additional criterion requiring that the neural response of one identity/category was at least 2 standard deviations (SD) above the mean of the neural responses of all others, which also allowed for identifying which identity/category the neuron’s responded to. These procedures are consistent with criteria employed in other related studies to detect responses to specific identities (De Falco et al., 2016; Rey et al., 2020).

After detecting neurons with responses to particular identities or categories, we employed a series of control analyses to further examine these neurons. We assessed the selectivity of each neuron to different identities for each neuron using an identity selectivity index defined as the *d’* between the most- and least-preferred identities (Grossman et al., 2019). This was computed as (μ_best_ - μ_worst_ / sqrt(0.5*(σ^2^_best_ + σ^2^_worst_))), wherein μ_best_, μ_worst_ and σ^2^_best_, σ^2^_worst_ denote the mean firing and variance of firing rate for the most- and least-preferred identities, respectively. We also computed a depth of selectivity (DOS) measure to summarize the responses of identity neurons, creating a scale that varies from 0, indicating equal responding to all identities, to 1 denoting exclusive responses to one identity but none of the others (Minxha et al., 2017; Wang et al., 2018). The DOS measure was quantified as (n-(sum(Rj / Rmax)/n-1), where n is the number of identities or categories (n = 50), Rj is the average firing rate to identity j, and Rmax is the maximum average firing rate across all identities. Finally, for each neuron, we also calculated the response ratio for each face/object identity. To do so, the responses of all identities were first divided by the response of the most preferred identity and then ranked from the most preferred to the least preferred, such that the response ratio of the most preferred identity is always 1. We compared response ratio for each ordered identity between identity vs. non-identity-selective neurons using two-tailed unpaired *t*-test (corrected for multiple comparisons using false discovery rate (FDR) (Benjamini & Hochberg, 1995). A steeper change from the best to the worst category indicates a stronger identity selectivity. Note that these measures were used to quantify the properties of identity neurons and compare them to non-identity neurons that were selected by the ANOVA procedure, and were not used to select identity neurons.

In the Treasure Hunt task, overall accuracy was evaluated based on the number of recall responses that were evaluated by the game as correct, based on the 13 virtual unit threshold. For the stimulus related responses, in Treasure Hunt, this relates to chest opening events, when stimuli are presented. We used t-tests to evaluate a significant change of firing, comparing the 1-second pre and post chest opening time, specifically for full chests, which contain presented stimuli.

To examine potential space- and sequence-related neural responses in TH, we analyzed the data from the navigation periods. For the space related analyses, we first binned the rectangular environment into a grid such that we could assign the position of the subject, the position of the targets (chests) and neural activity as relating to particular bins. This allowed us to examine relationships between neural firing and the subject’s and/or target’s positions. In doing so, we used a minimum occupancy of 1 second, and excluded stationary periods (if the speed was lower than 5×10^−6^ virtual units / second) for a bin to be included in any subsequent analyses. After these exclusions, we computed the average firing rate in each grid by binning the associated spikes based on the position of the player or of the targets respectively. For place cell analyses, a spatial binning of 5-by-7 across the navigable range was used to associate spiking with the players position. For target cell analyses, a binning of 2-by-4 across the possible chest range was used to associate spiking with the target destination. One-way ANOVAs were used to evaluate whether firing rates were significantly modulated by subject or target spatial location, assessed using the aforementioned surrogate procedure. For visualization purposes, we smoothed the firing rate heatmaps by binning into a finer 5-by-8 grid, with a Gaussian filter with a 1.1-bin SD. We additionally tested for a modulation of firing rate by serial position, using an ANOVA procedure to evaluate whether each neuron’s firing rate during navigation was significantly modulated by the serial position of the four chests that were presented per trial.

After analyzing cell responses within each task – including stimulus-related responses in both tasks, identity and object category responses in the one-back task, and place, target and serial position related responses in the Treasure Hunt task – we next tested for interactions betweens neuronal representations across tasks. Specifically, we tested for an over-representation of neurons with representation A in the one-back task and representation B in the Treasure Hunt task, across all analyzed representations ‘A’ and ‘B’. We did so by computing the number of neurons that overlapped across each task, as related to the number of neurons with that representation within each task, and evaluated the statistical significance of the overlap with a chi-square test. This procedure allowed for evaluating whether there was an over-representation of neurons with a particular representation in one task and a similar or different representation in the other task. As well as comparing the number of identified neurons, we also examined the full set of statistical test values for each analysis (either t-values of f-values, depending on the analysis), and compared them between analyses. For each combination of task responses, we visualized the relationship between the statistical values and computed the spearman rank correlation between them.

To visualize potential relationships between responses to different features, we did a series of simulations. Each simulation consisted of bivariate distribution, representing responses to ‘task 1’ and ‘task 2’, with 500 values per task, simulated with different relationships between simulated task responses. To simulate correlated responses, we simulated bivariate normal data with either a correlation of r = 0.85 (correlated example) or r = 0 (uncorrelated example). To simulate data with task specific responses, univariate normal data was first independently simulated for each task response, and then combined such that individual data points responded to one or the other task, split evenly. Finally, to simulate a combination of task-specific and task-general responses, we combined 80% of data points sampled as task specific responses with 20% sampled as correlated responses. Each of these simulated cases creates distributions with idiosyncratic properties that can be used to visually compare to the observed set of responses in the empirical data. The full set of parameters used to simulate the data is available in the project repository.

## Results

In this project, we used a paired-task session (Fig 1A), in which 5 neurosurgical patients completed 22 paired task sessions consisting of a working memory one-back (OB) task followed by the Treasure Hunt (TH) spatial navigation task (Fig 1A&B). There were two versions of these paired-task sessions, one using face stimuli and the other using object stimuli. We recorded from implanted microwires during the paired task session, that were pre-processed together such that the same single-neurons could be compared across both tasks (Table 1; face-version: 608 neurons, object-version: 565 neurons), across the hippocampus and amygdala (Fig 1D). Within each task, single-neuron responses were analyzed based on previous analyses of these tasks, finding stimulus-related and identity related responses in the one-back task (Cao et al., 2021; Cao, Wang, et al., 2022) and space and sequence related responses in the Treasure Hunt task (Tsitsiklis et al., 2020). We then compared responses across the two tasks, identifying neurons that have task specific responses, neurons that maintain a representation across task contexts, and neurons that switch representations between tasks.

**Figure 1.**
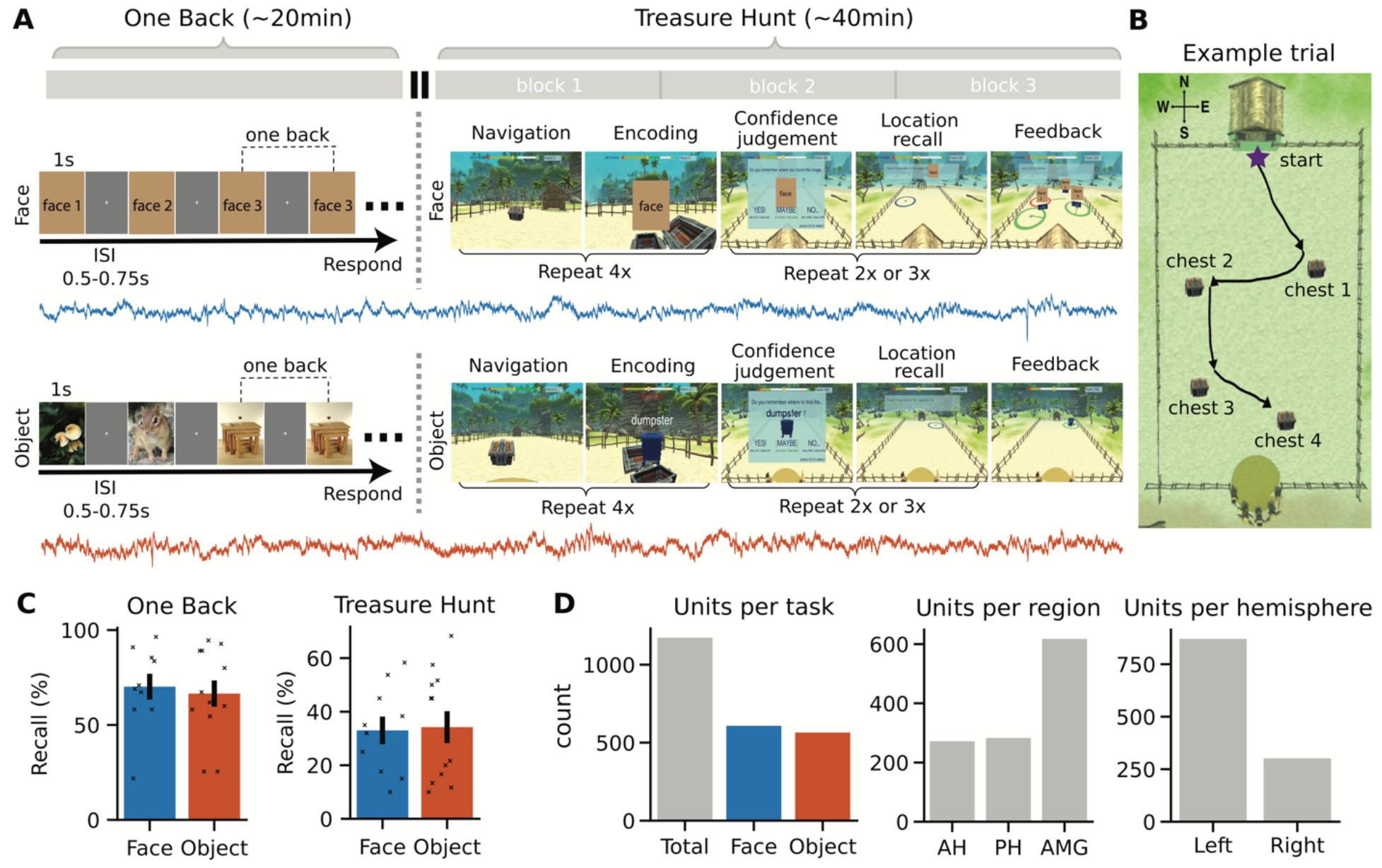
Paired-task session with combined spike sorting. A) Overview of the paired task session including a one-back task (left) and Treasure Hunt task (right), each of which had versions with either face (top; blue) or object (bottom; orange) stimuli. Here, faces have been replaced with place-holders – note that in the presented version, the task showed real pictures. The time trace represents the neural recordings that were recorded across both tasks in a combined session. B) A top down representation of the Treasure Hunt arena, showing the layout and an example trial with 4 chests. Note that this view is not shown to participants, and during gameplay each chest is presented serially. C) Behavioral performance for the one-back (left) and treasure hunt (right) tasks. D) Number of identified neurons across stimulus variants (left), anatomical region (middle), and hemisphere (right).

### 3.1 One-Back Results

Behaviorally, participants performed consistently well at the n-back task, both in terms of accuracy of repetition detection (Fig 1C; OB-face: 70.18% ± 21.52 [mean ± 1 standard deviation across sessions]; OB-object: 66.52% ± 23.98), and reaction time (OB-face: 203.7 ± 145.4ms; OB-object: 118.7 ± 101.1 ms). Comparing between stimulus version, there was no significant difference in accuracy (t(20)=0.38, p=0.71, unpaired two-sample *t*-test) or reaction time (t(20)=1.61, p=0.12, unpaired two-sample *t*-test) between the face and object versions of the one-back task. Analyzing the neural data, we first examined whether there were stimulus reactive cells with a significant activation for presented images, using paired t-tests (Fig 2A&D). We detected a significant number of stimulus-responsive cells across both task variants (Fig 2G; OB-face: 102/608 (16.78%), p<10^−25^, one-sided binomial test; OB-object: 69/565 (12.21%), p<10^−10^, one-sided binomial test).

**Figure 2.**
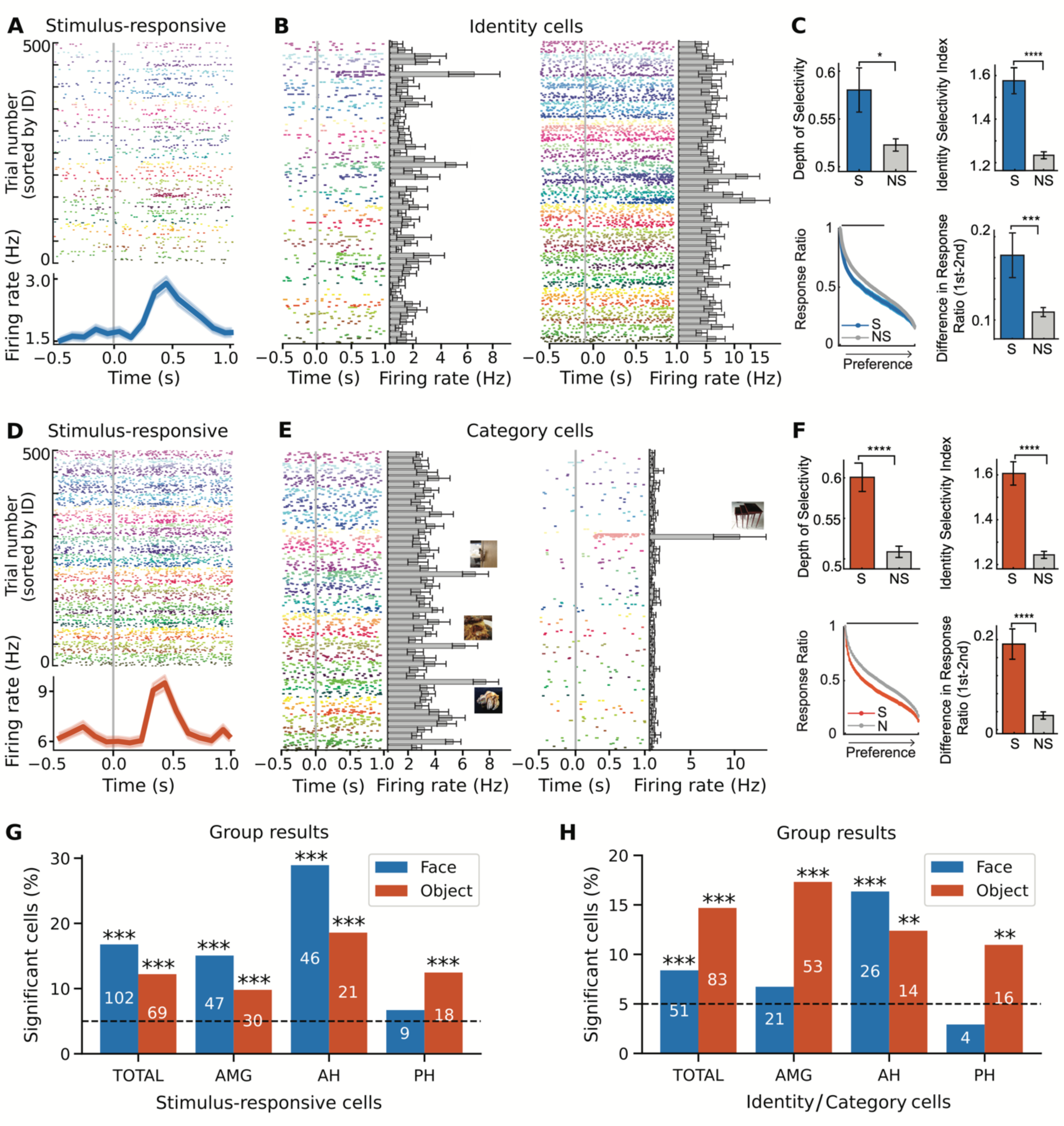
Individual neurons in the one-back task respond to stimuli, individual identities, and object categories. A-F) Responses in the one-back task, for the face (A-C) and object (D-F) versions. Raster plots show neuronal responses to 500 faces (A, B) or objects (D, E). Each identity is coded by different colors. Trials are aligned to the stimulus onset (gray line) and are grouped by individual identity. A&D) Example neurons showing a stimulus related increase in activity for face (A) and object (D) stimuli. B&E) Example neurons with responses to specific identities (B) or object categories (E). Bar plots show average response for each identity and error bars denote ±SEM across face or object examples. Sample stimulus from encoded identities is displayed on top of the bars (face images have been removed). C&F) Group analyses of identity/category-selective neurons (S: selective; N: non-selective), including depth of selectivity (DOS; top left), identity selectivity index (top-right), response ratio (bottom left), and difference in response ratio between the first and second most preferred identities (bottom-right). The response ratio plot shows ordered average responses from the most- to the least-preferred identity, normalized by the response to the most-preferred identity. The black bar refers to the significant difference between identity-selective vs. non-selective neurons (two-tailed unpaired t-test, p<0.05, corrected by FDR for Q < 0.05). Asterisks indicate a significant difference under two-tailed unpaired t-test, *: P < 0.05, ** P < 0.01, *** P<0.001, **** P < 0.0001. Error bars denote ±SEM across neurons. G&H) Group-level results, showing the number of significant cells in the face (blue) and object (orange) versions of the task, as well as across regions (AMG: amygdala; AH: anterior hippocampus; PH: posterior hippocampus).

We subsequently analyzed neural responses for cells that responded selectively to individual identities (people or objects) using 1-way ANOVA (see methods for details; Fig 2B&E). We found a significant number of identity cells in both task variants (Fig 2H; OB-face: 51/608 (8.39%), p=0.0003, onesided binomial test; OB-object: 83/565 (14.69%), p<10^−17^, one-sided binomial test). To further examine the identity neurons, we compared selective and non-selective cells on several measures, including a depth of selectivity index, an identity selectivity index, and on differences in response ratios (see methods). As compared to non-selective neurons, identity-neurons had a significantly higher depth of selectivity index, a significantly higher identity selectivity index, and a significantly higher difference in response ratios between the first and second most-preferred identities (Fig 2C&F; all p’s<0.05, two-tailed unpaired t-test), indicating that their response was selective to specific face or object identities.

### 3.2 Treasure Hunt Results

In the Treasure Hunt task, we measured performance on each trial based on the distance between the subject’s response location to the item’s actual position. Participants responded accurately on ~33% of trials (Fig 1C; TH-face: 33.0% ± 15.50 [mean ± 1 standard deviation across sessions]; TH-object: 34.2% ± 19.77). There was no significant difference in performance between that face and object versions of TH (t(20)=-0.15, p=0.88, unpaired two-sample *t*-test). Analyzing the neural data, we first tested for stimulus responsive cells during the chest-opening events (Fig 3A&D), finding a significant number of stimulus-responsive cells (Fig 3G; TH-face: 108/608 (17.76%), p<10^−29^, one-sided binomial test; TH-object: 61/565 (10.80%), p<10^−7^, one-sided binomial test).

**Figure 3.**
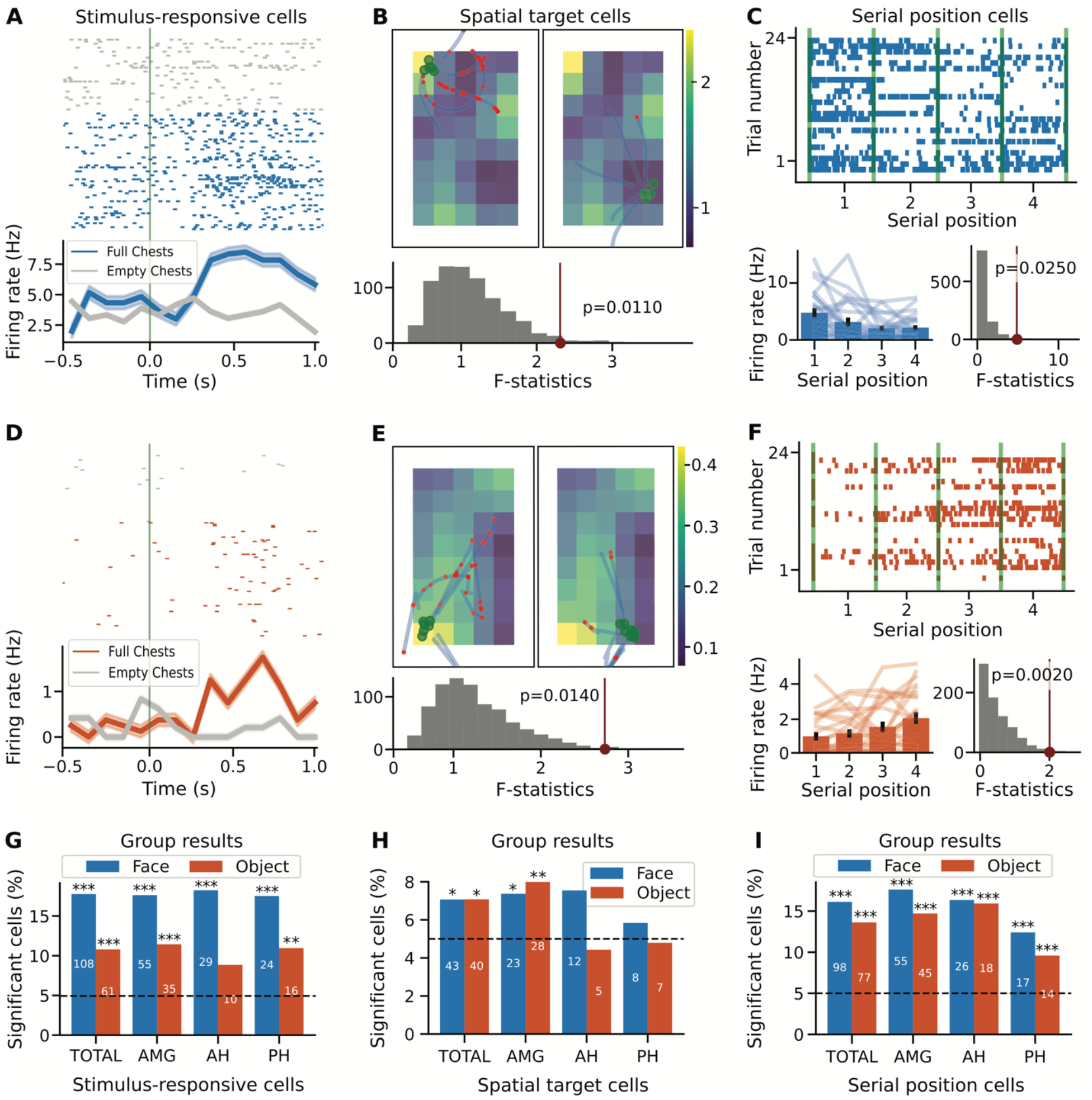
Individual neurons in the Treasure Hunt task respond to stimuli, target location, and serial position. Neurons in the Treasure Hunt task were identified if they responded to stimulus presentations (A,D,F), spatial features (B,E,H), and/or serial position of presented items (C,F,I). For all analyses, the first two rows show example neurons (top / blue: face version; bottom / orange: object version) and the bottom row shows group level results, across stimulus variants and regions. A&D) stimulus responsive neurons were identified if they have a significant response to presented stimuli. B&E) spatial target cells were identified based on having a firing field related to the location of the target (chest) location. C&F) Serial position cells were identified based on having a differential firing rate related to the serial position of the four chests presented per trial. G-I) Group-level results, showing the number of significant neurons in the face (blue) and object (orange) versions of the task, as well as across regions (AMG: amygdala; AH: anterior hippocampus; PH: posterior hippocampus).

We next looked for space-related representations, including firing patterns that related to the subject’s location as well as spatial target locations. We first looked for place cells, by examining firing rates in relation to the player’s position in virtual space, and although there were some individual neurons that passed the statistical test, overall there was not a significant number of place cells in this task (TH-face: 28/608 (4.61%), one-sided binomial test p=0.6983; TH-object 28/565 (4.96%), one-sided binomial test p=0.5462). When looking for spatial target cells, by examining firing rates in relation to the player’s target destination (Fig 3B&E), we did find a significant number of spatial target cells (Fig 3H; TH-face: 43/608 (7.07%), one-sided binomial test p=0.0155; TH-object: 40/565 (7.08%), one-sided binomial test p=0.0187). We additionally analyzed the Treasure Hunt data for serial position representations, as shown in previous datasets, by examining whether there was a modulation of firing activity during navigation based on the serial order of the four chests that were presented per trial (Fig 3C&F). We found a significant number of serial position cells in both variants of the Treasure Hunt task (Fig 3I; TH-face: 98/608 (16.78%), p<10^−23^, one-sided binomial test; TH-object: 77/565 (13.63%), p<10^−14^, one-sided binomial test).

### 3.3 Overlap Results

We next examined the overlap between neuronal responses across the two tasks, starting by comparing stimulus responses across both tasks. To do so, we used Chi-squared tests to examine the number of cells that responded across one or both tasks. We found a significant overlap between the neurons that were found to be stimulus-responsive in one-back and those that were stimulus-responsive in Treasure Hunt (Fig 4A-C; face-version: 31 overlap neurons (30.39%), p=0.0003, Chi-squared test; Fig 4D-F; object-version: 15 overlap neurons (21.74%), p=0.0018, Chi-squared test). Similarly, there was evidence of an over-representation of identity representation in the one-back task, when compared with more general stimulus response in Treasure Hunt, though this was only trending in the face version (face-version: 14 overlap neurons (27.45%), p=0.0586, Chi-squared test; object-version: 15 overlap neurons (18.07%), p=0.0207, Chi-squared test). This overlap suggests that a significant number of neurons maintained a consistent representation across tasks.

**Figure 4.**
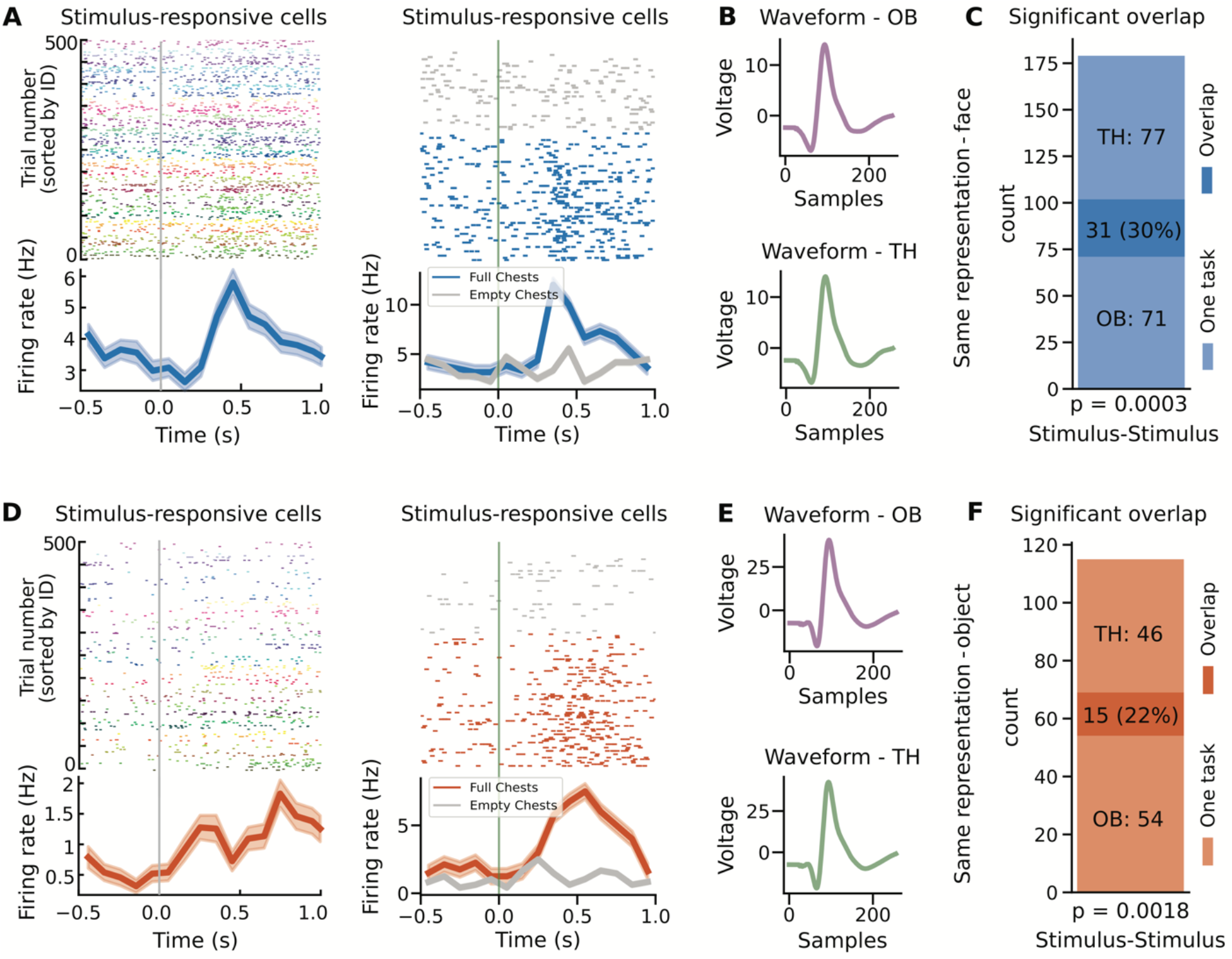
Overlap results for same representation. Examples of neurons with responses across both task contexts for neurons with the same kind of representation, responding to stimuli presentations across both tasks, for face stimuli (A-C) and for object stimuli (D-F). A&D) Example neurons that respond to stimuli in both the one-back and Treasure Hunt task. B&E) Average waveforms for the example neurons in A&D split across tasks. The waveforms are very similar, motivating that these detected units are well isolated and represent the same neuron across tasks. C&F) Shows the number of task responsive neurons, including the number of neurons that respond in only one task, and the number of overlap neurons that respond in both tasks.

We then measured the cells that shifted the nature of their representations between tasks, examining the stimulus- and identity-responsive cells from the one-back task as compared to target- and serial-position-responsive cells from the Treasure Hunt task. We found that that neurons that were stimulus responsive in the one-back task often responded at particular serial positions in Treasure Hunt, though this overlap was only significantly over-represented with the object stimuli (Fig 5A-C; face-version: 18 overlap neurons (17.65%), p=0.6453, Chi-squared test; Fig 5D-F; object-version: 17 overlap neurons (24.64%) p=0.0044, Chi-squared test). This pattern of results, in which the neurons that respond to all stimuli during the one-back task shift to represent particular serial positions during Treasure Hunt, provides an example of how some neurons can change the nature of their coding fundamentally between different behavioral task settings. The full set of overlap results, including non-significant overlaps, is available in Table 2.

**Figure 5.**
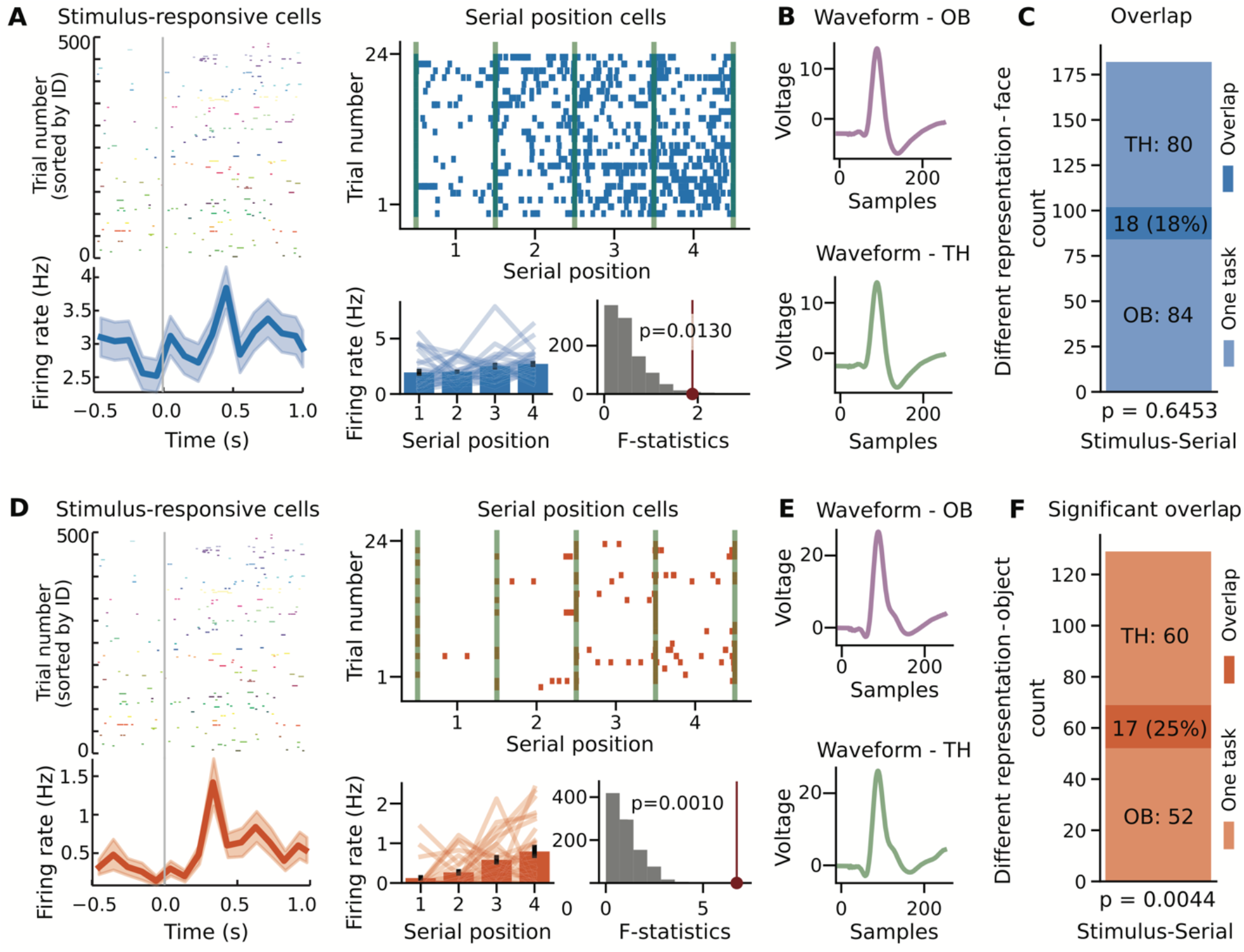
Overlap results for different representations. Examples of neurons with responses across both task contexts, for neurons with different representations, responding to different aspects of each task for face stimuli (A-C) and for object stimuli (D-F). A&D) Example neurons that respond to stimuli in the one-back task and to serial position in the Treasure Hunt task. B&E) Average waveforms for the example neurons in A&D, split across tasks. The waveforms are very similar, motivating that these detected units are well isolated and represent the same neuron across tasks. C&F) Shows the number of task responsive neurons, including the number of neurons that respond in only one task, and the number of overlap neurons that respond in both tasks.

**Table 2.** Full set of overlap results. The full set of comparisons between tasks. Reported p-values are for Chi-squared tests evaluating whether the number of neurons that overlap across tasks is different than expected by chance.

| OB cell labels | TH cell labels | Number of cells in OB | Number of cells in TH | Number of cells in overlap | Percentage of cells overlap | p-value |
| --- | --- | --- | --- | --- | --- | --- |
| Stimulus | Stimulus | 102, 69 | 108, 61 | 31, 15 | 10.39%, 21.74% | 0.0003, 0.0018 |
|  | Spatial target |  | 43, 40 | 7, 7 | 6.86%, 10.14% | 0.9279, 0.2894 |
|  | Serial position |  | 98, 77 | 18, 17 | 17.65%, 20.64% | 0.6453, 0.0044 |
| Identity/Category | Stimulus | 51, 83 | 108, 61 | 14, 15 | 27.45%, 18.07% | 0.0586, 0.0207 |
|  | Spatial target |  | 43, 44 | 2, 8 | 3.92%, 9.64% | 0.3591, 0.3251 |
|  | Serial position |  | 98, 77 | 9, 13 | 17.65%, 15.66% | 0.7564, 0.5586 |

Finally, to examine the full pattern of results across all neurons, for task combinations for which we observed a significant number of overlap neurons, we examined the pattern of statistical measures (t-values or f-values) for each analysis. In order to illustrate different potential patterns, we used simulated data to demonstrate four possible relationships of responses between tasks, including having correlated responses, having uncorrelated responses, having task specific responses, and having a combination of task specific responses and neurons that respond to both tasks (Fig 6A-D). The empirical distributions appeared qualitatively consistent with the simulated data that was created with a combination of task-specific and overlapping responses (Fig 6E-F). Specifically, in the empirical data, there were no significant correlations between the statistical measures (spearman correlation, all p’s > 0.05), which is also consistent with their being a combination of task specific and task general responses. Overall, we conclude from these analyses that there are neurons that do respond across tasks, but that this appears to represent a subset of neurons, as many neurons appear to respond in a task-specific manner.

**Figure 6.**
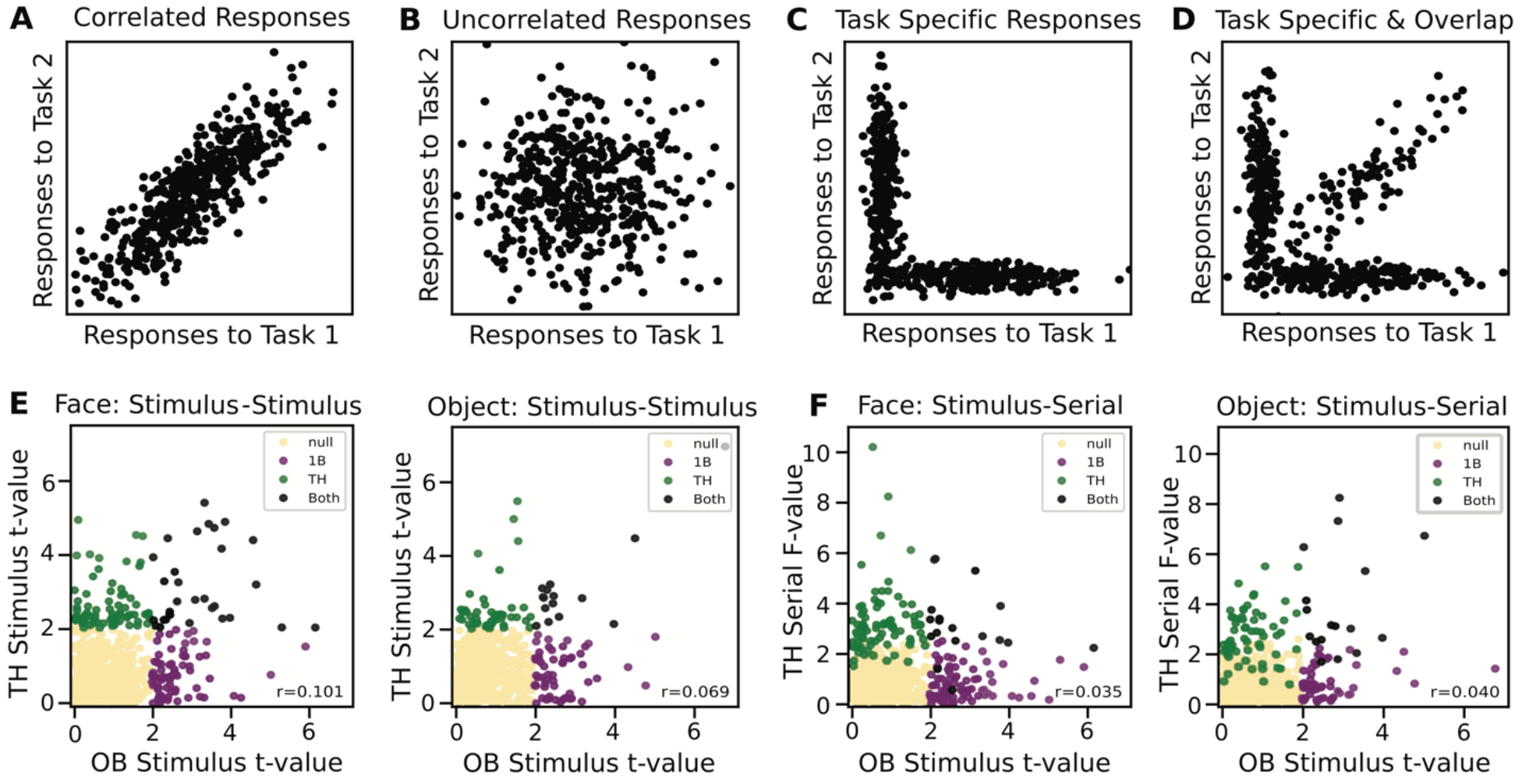
Group level comparison of responses across tasks across all neurons. A-D) Simulated data showing hypotheses of potential relationships of the task-related activity across two distinct tasks. Four potential relationships are shown: A) responses are correlated between tasks, B) responses are uncorrelated across tasks, C) responses are task-specific, such that individual neurons respond to single task only, D) there is a combination of task-specific responses, whereby most neurons respond to one task only, with a subset of neurons that respond to both (‘overlap’ neurons). E-F) Empirical distributions comparing the statistical measures for different responses between tasks, shown for the pairings in which we saw evidence of a significant number of overlap neurons. Each data point represents an individual neuron, plotted based on the statistical measures computed for different analyses (f-value or t-value, depending on the analysis – note that t-values are absolute valued). Each data point is colored by the outcomes of the analyses (yellow: not significant in either task; purple: only significant in the one-back task measure; green: only significant in the Treasure Hunt task measure; black: significant in both the one-back and Treasure Hunt task measures. Inset text shows the correlation values. The overall pattern of the empirical data is most consistent with the simulated hypothesis in which there is a combination of task specific and overlap neurons.

## Discussion

In this study, we leveraged the human capacity for rapidly shifting task demands to investigate representations in individual neurons in the MTL across distinct behavioral contexts. We did so using a paired-task session including a one-back visual working-memory task followed by a spatial-navigation and memory task with human participants. Of the neurons that responded in both tasks, some neurons do maintain a consistent representation, by responding to the same class of stimuli across task contexts, whereas other neurons appear to switch their representation, responding to one type of feature in one task, and switching to represent a different, seemingly unrelated, feature in a subsequent task. Our findings thus contribute to the discussion on the specificity (or lack thereof) of neural representations at the level of individual neurons. Relatedly, it helps to address an implicit tension in the literature – given an ever expanding number of experiments that often report between ~5-50% of neurons that respond to a particular variable of interest, it quickly becomes practically impossible that each of these reflects a unique and independent set of neurons with a consistent feature encoding. Addressing this tension, here we show that while individual neurons in the human MTL can have what appear to be specific and selective responses within a particular context, across different tasks with disparate demands, neurons can flexibly shift between different representations.

We note that while this experiment motivates that individual neurons can have a variable representation across contexts in the MTL, this is distinct from the notion of ‘mixed selectivity’ in which individual neurons respond to multiple features simultaneously. For example, in the prefrontal cortex, individual neurons with ‘mixed selectivity’ can non-linearly encode combinations of multiple features, suggesting a code in which variables of interest are thought to be represented across ensembles of neurons (Duncan, 2001; Fusi et al., 2016; Rigotti et al., 2013). The findings in our study are distinct from this notion of broadly tuned neural responses, and do not suggest that MTL neurons respond broadly to a mixture of features within the same task, but rather that they can have a selective response in one context, which flexibly reorganizes to a different selective response in another context. This distinction emphasizes that there is likely a high degree of regional specificity in the nature of neural representations, with different brain networks using distinct strategies to encode task-relevant information.

The particular design and task set used in this experiment also reflect a targeted combination of MTL related functions, including having representations related to concepts (Quiroga, 2012) and to spatial navigation (Moser et al., 2017). The combination of tasks in this experiment were specifically tuned to relate to these seemingly distinct functions of the MTL, with much discussion about the relation of these distinct functions, and how they each relate to memory. We found that neurons in the same regions engage in both spatial and concept-related tasks, with a clear overlap between cells engaged in the two tasks, replicating and extending previous studies using each in isolation to show spatial (Tsitsiklis et al., 2020) and identity-related responses (Cao et al., 2021; Cao, Wang, et al., 2022) in the human MTL. We also ran these tasks with two distinct sets of stimuli (faces and objects), and while there were some small differences, there was not overall a clear distinction between the two different stimulus versions of the tasks.

One way to contextualize our findings is in relation to the notion of ‘cognitive maps’ (Behrens et al., 2018; Schiller et al., 2015; Tolman, 1948), or similarly considering the MTL as a relational processing system (Eichenbaum & Cohen, 2014). Under this framework, the hippocampus and surrounding areas can be thought to represent abstract relations across dimensions, whereby space is a prominent, but non-exclusive, domain for which a map can be constructed. Our experiment is consistent with the MTL being involved in generalized relational representation, finding responses that relate to space, sequences, and stimulus categories within a single recording session. We note however, that while our experiment is consistent with the general notion of ‘cognitive maps’, the specific patterns of changing representations are not clearly the pattern of dynamic representation that might be predicted under this framework. It could be hypothesized, for example, that individual neurons in a ‘cognitive map’ would have a conserved function of representing locations in different dimensional spaces across contexts, representing for example, locations in physical space in one task context and location in stimulus space in another. The changes in representations we observed were largely related to more broadly tuned representations such as stimulus responses, which are not entirely consistent with such a change in representation that relates to mapping specific locations in different feature spaces. Future work should seek to continue to evaluate individual neurons across distinct tasks in order to better evaluate how and understand why neurons appear to change their representations.

This experiment emphasizes that the MTL is able to flexibly adapt to representing task-relevant features across changing behavioral demands. This may also relate to a potentially surprising aspect of our results, whereby we found a significant number of spatial target cells, but that we did not find a significant level of place cells. This general finding, in which there is stronger evidence for spatial target encoding rather than player location, is consistent with previous analyses of an independent dataset using this task (Tsitsiklis et al., 2020). This kind of encoding of remote target locations is reminiscent of non-human primate work in which MTL neurons often represent remote locations (Rolls & Wirth, 2018), including spatial-view cells which encode the viewing location of the monkey (Rolls, 1999), grid-like representation of visual space (Killian et al., 2012), and schema cells which respond to abstract representations of space (Baraduc et al., 2019). Viewpoint-specific representations of visual scenes have also been observed in human MTL with fMRI (Epstein et al., 2003). In contrast to our findings, other human work using different virtual-navigation tasks has found place cells (Ekstrom et al., 2003; Miller et al., 2013). We thus hypothesize that the predominance of spatial-target responses observed here may relate to the behavioral demands. Notably, the Treasure Hunt task has an emphasis on the remote location of visible chest locations, rather than on current location, such that participants likely focused on the remote target locations while navigating, which is reflected in the spatial target cells. This shift in the nature of location tuning between place and spatial-target cells is consistent with the broader pattern we found here between the one-back and spatial tasks, where human MTL neurons flexibly vary their coding properties depending on task demands.

Beyond single-neuron recordings, our findings are also consistent with other human work that has emphasized that the same circuits seem to be involved in multiple distinct feature representations and tasks. Based on a review of human studies predominantly using fMRI, the human MTL has been found to have a similar functional organization in relation to spatial navigation as has been established in animal models, including elements of the hippocampal network that activate during specific spatial settings (Epstein et al., 2017). Building on this homology, other fMRI experiments have established that the same or similar circuits are engaged when participants engage in non-spatial tasks that can still be conceptualized as ‘navigating’ feature spaces. For example, in an experiment in which participants made social decisions in a role playing game, analyzing the task decisions as movements through a social space revealed task related activations in the hippocampus (Tavares et al., 2015). In another experiment that involved making decisions about items varying across a 2d feature space, grid-like activations were found in a set of regions that are also related to spatial navigation (Constantinescu et al., 2016). While these fMRI studies do not allow for inference on the activity of single neurons, they reflect a growing literature that is establishing that the same MTL circuits engage in different tasks, including across the domains of spatial tasks, faces, and object categories, within which our current work helps to evaluate the single-neuron correlates of how these circuits become flexibly recruited across task contexts.

A key limitation of the previous literature as it pertains to explaining the relations of representations across contexts is that the majority of previous literature examines how individual neurons represent variables of interest in experiments using only a single task or across a limited behavioral context. One reason for this is that most animal models require extensive training on behavioral tasks, and it is therefore practically difficult to have animals quickly and flexibly maneuver between completely different tasks while recording across the same neurons. Here, we leveraged the benefits of working with humans, which allows for rapidly switching between multiple complex tasks. By using a paired-task session, this research design required only minimal practical updates to deploy a protocol in which two distinct tasks could be run adjacent in time in order for the data to be spike sorted and analyzed together. In future work, this experimental design could continue to be applied in other human single-neuron recording settings across different task combinations in different brain areas to further probe questions about neural representations across task contexts. Although some of these questions can also be examined with fMRI, which conveniently allows for multiple scans in the same individuals, single-neuron recordings have the advantage of providing much higher spatial and temporal precision that is useful for measuring precisely timed neuronal signals such as those that represent specific locations during active navigation.

We also note that it is typical for human single-neuron research participants to complete multiple different tasks across the 1-2 weeks that they are typically in the EMU. In principle, the kinds of comparisons performed in this study could be extended across more tasks, potentially even with retrospective data, pending some technical challenges of aligning and analyzing data across recording sessions done at different times. Most notably, this requires having strategies for aligning spike sorting solutions across recording sessions in order to align putative single units. While it is unclear to what extent historical data may be alignable, prospectively recording long-term collections of ongoing data may also provide a strategy for analyzing data across sessions (Chaure & Rey, 2020), though more work is needed with this approach. Overall, we propose that analyzing single-neuron activity across task contexts is a fruitful research strategy that leverages the benefits of working with humans in order to answer important questions about neural representations.

## Conclusion

The medial temporal lobe is a complex structure that is known to be involved in multiple cognitive processes, including spatial navigation, representing high-level concepts, and memory processing. Here, we investigated the relation between these seemingly distinct functions, by using a paired task session in which the activity of the same neurons can be evaluated across task contexts. By doing so, we were able to show that there are multiple patterns of neural activity across tasks, whereby some neurons are active only in one task context, some neurons maintain a similar representation, and some neurons appear to switch their representation entirely, responding to seemingly distinct features in different task contexts. By showing that individual neurons can change the nature of their coding scheme between different behavioral tasks, our results contribute to a broader understanding of how the brain supports a broad range of behaviors. Our results show that individual neurons change their coding scheme between behaviors, demonstrating that, at least in the medial temporal lobe, there is a substantial degree of flexibility in neural networks, as opposed to requiring dedicated brain regions for individual behavioral tasks and types of neural representations. This topic may be a useful area for future research that could assess the modularity and flexibility of neural coding more generally and characterize the principles that explain the transformations in neural coding by individual neurons that appear between tasks.

## Acknowledgements

We would like to thank all patients for their participation, and staff from WVU Ruby Memorial Hospital for support with patient testing.

## Abbreviations

MTL: medial temporal lobe
iEEG: intracranial electroencephalography
NWB: Neurodata Without Borders
EMU: epilepsy monitoring unit

## Materials Descriptions & Availability Statements

### Project Repository

This project is openly available through an online project repository, which includes all the code used for data collection and analysis, as well as step-by-step guides through the analyses.

Project Repository: https://github.com/HSUpipeline/AnalyzeTH

### Datasets

This project uses electrophysiological data collected from neuro-surgical patients. The data from the working memory task is openly available as part of a larger dataset (https://osf.io/824s7/). The data from the spatial task will be made available prior to publication. All subject testing and data processing complied with the policies of local institutional review boards.

### Software

Code used and written for this project was written in the Python programming language. All the code used within this project is deposited in the project repository and is made openly available and licensed for reuse.

Management of the dataset was done using the Human Single Unit (HSU) Pipeline:

HSU pipeline: https://github.com/HSUPipeline

Analyses of the single-neuron data were done using the spiketools toolbox:

spiketools repository: https://github.com/spiketools/spiketools

---

**Supplemental Figure 1.**
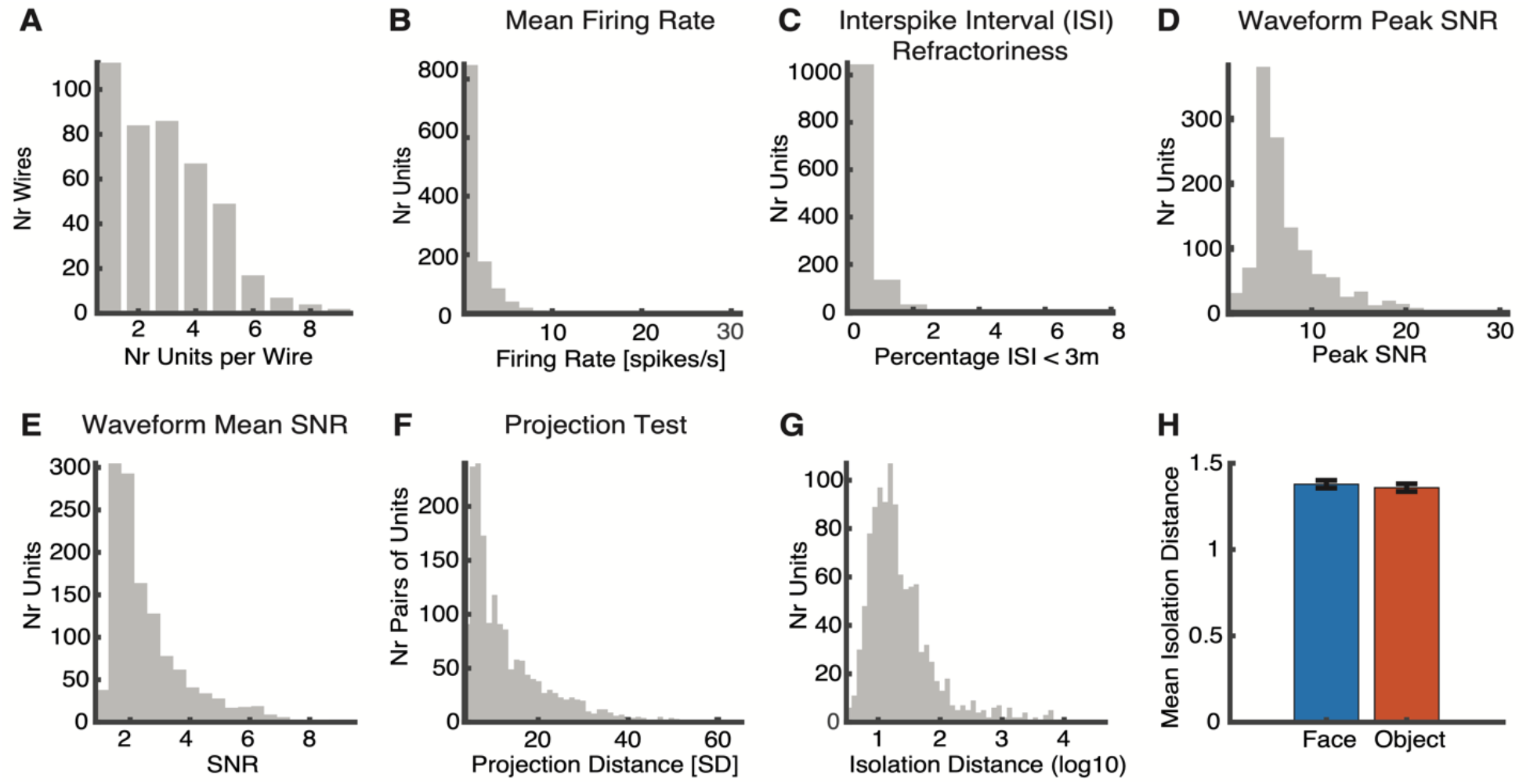
Spike sorting quality measures. Panels A-F reflect spike sorting metrics combined across the face and object versions of the task, including A) the number of units per wire, B) the firing rates, C) the inter-spike intervals, D) the waveform peak signal-to-noise ratio (SNR), E) the waveform mean SNR, F) the projection distance, and G) the isolation distance. G) The mean isolation distance averaged across face and object versions of the paired task sessions.

**Supplemental Figure 2.**
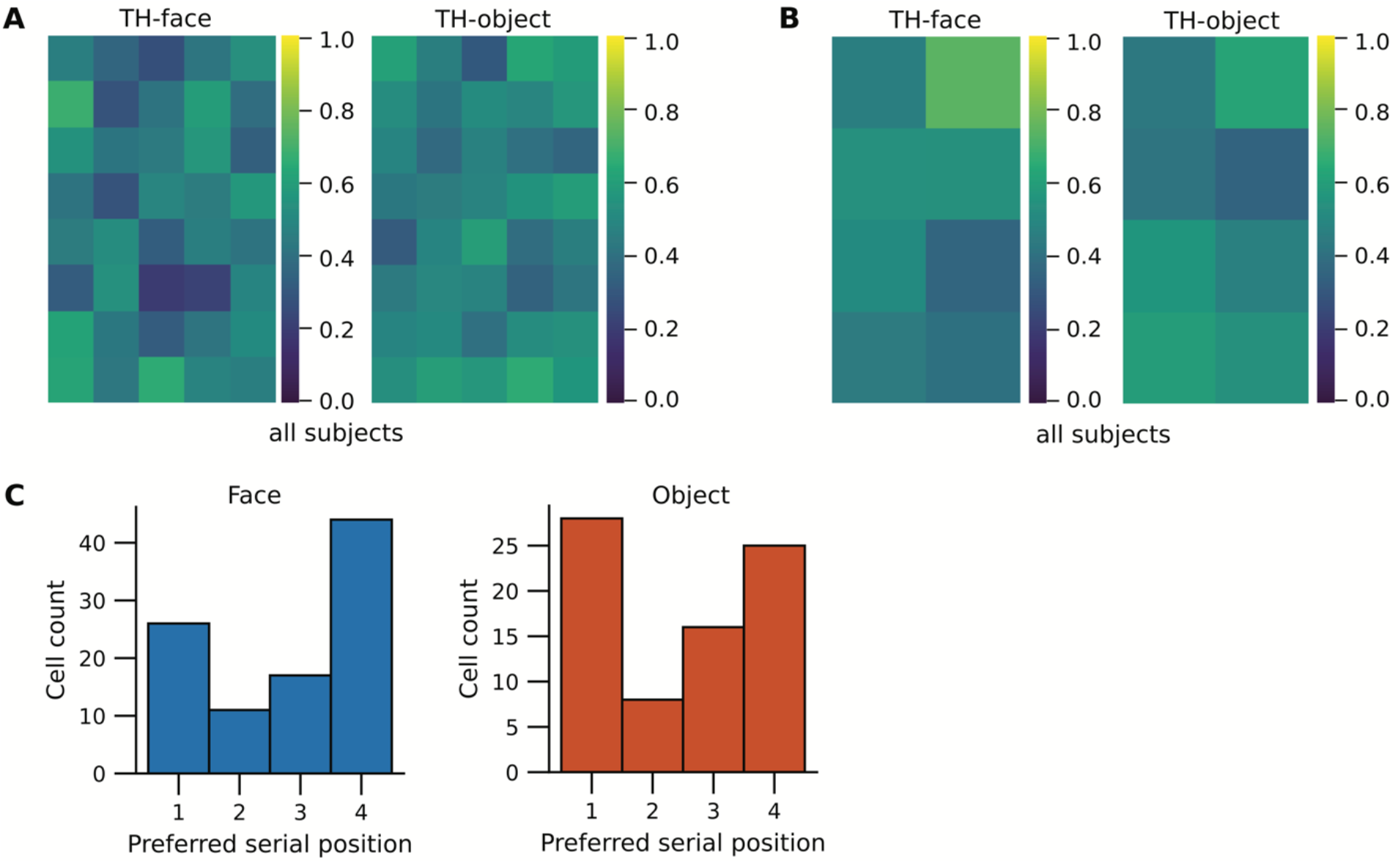
Group-level analyses of the Treasure Hunt task. A) Averaged firing rate maps for all identified place cells, split by face and object versions of the task. B) Averaged firing rate maps from all significant spatial target cells, split by face and object versions of the task. C) The count of preferred positions for all significant serial-position cells, split by face and object versions of the task.

